# Awake fMRI Reveals Covert Arousal in Aggressive Dogs Under Social Resource Threat

**DOI:** 10.1101/203323

**Authors:** Peter Cook, Ashley Prichard, Mark Spivak, Gregory S. Berns

## Abstract

Domestic dogs are highly social, and have been shown sensitive not only to the actions of humans and other dogs but to the interactions between them. To examine the canine neurobiological response to observed interactions between a human and another dog, we collected fMRI data from dogs while they watched their owner feed a realistic fake dog or deposit food in a bucket. Given the likelyihood that arousal and affective state may contribute to responses to observed social situations, we examined the relationship between amygdala activation in these two conditions and an independent measure of aggressive temperament from the C-BARQ scale. Dogs rated more aggressive showed significantly higher activation in the fake-dog versus bucket condition. This finding suggests a neurobiological mechanism mediated by the amygdala for dog-directed aggression, especially when their owner interacts with another dog. Such a mechanism may have some parallels to human jealousy. Further, it adds to a growing body of evidence that specific neurobiological responses correlate with canine temperament and can be a predictor of future behavior. We also found evidence that the amygdala response habituates with repeated observed interactions. This suggests value in exposure-based interventions for potentially aggressive dogs.

## Introduction

Social relationships with human caregivers are central to domestic dog behavioral ecology. Most dogs rely on humans for food, shelter, and companionship, and extensive evidence shows that dogs are attentive to and sensitive to human social signals, including tone of voice, posture, and facial expression (Topál *et al.*, 1998; Schwab & Huber, 2006; Palmer and Custance, 2008; Lakatos et al., 2012; Merola et al., 2012; Müller *et al.*, 2015; Nagasawa et al., 2015; Müller et al., 2015). Given the centrality of this relationship to a dog’s life, it is perhaps not surprising that dogs might be aware of potential challenges to their privileged social access to familiar humans, and even act to protect that access. Recent behavioral studyies have shown that some dogs act aggressively toward a fake dog their owner praises, but not toward socially irrelevant inanimate objects their owner praises (Harris & Prouvost, 2014). In addition, similar to primates (Brosnan & De Waal, 2003), dogs are sensitive to inequity in reward, and react negatively when another dog receives a preferred reward (Range et al., 2009; Range, Leitner & Viranyi, 2012). In combination with the reported prevalence of dog-dog and dog-human aggression in social situations involving interactions between a caregiver and another dog or human (Wright, 1991; Casey et al., 2014), this suggests that a form of social resource guarding may contribute to aggressive behavior in domestic dogs. This type of socio-emotionally driven behavior might be interpreted as ‘jealousy,’ as has been explored previously in primates (Cubicciotti & Mason, 1978; Rilling, Winslow & Kilts, 2004; Panksepp, 2010). Nevertheless, regardless of the interpretation of the behavior, better understanding of the neurobiological phenomena underlying such canine social resource guarding might explain the proximate cause of competitive aggression. Such understanding might be of use in predicting which dogs are more likely to be aggressive, in what situation(s), and what behavior modification or preventive management strategies might be most effective when addressing aggressive proclivities.

In previous work (Berns, Brooks & Spivak, 2012; see reviews: Cook et al., 2016; Berns & Cook, 2016), we have used unrestrained, awake-fMRI with domestic dogs to examine the neurobiological underpinnings of social behavior and their relationship to canine temperament. Because we train subjects with operant conditioning and positive reinforcement, and participation is always voluntary, this allows us to perform human-style cognitive neuroscience with dogs, assessing brain response in alert animals in a range of affective states. For example, we have found that brain responses can be strong predictors of future behavior. In a recent study, we found that dogs who showed greater striatal activation for expectation of verbal praise, relative to food, were just as likely to show a behavioral preference for owner interaction versus opportunity to eat (Cook et al., 2016). We have also found that temperament as assessed by owner relates to degree of activation in striatal reward learning regions in response to learned signals from familiar and unfamiliar humans (Cook, Spivak & Berns, 2014). In addition, in a study featuring 50 dogs undergoing training for service work, amygdala and striatal activation patterns to learned signals were strong predictors of placement success (Berns et al., 2017). In certain cases, where behavioral response may be affected by a wide range of uncontrolled factors, or is unlikely to emerge except in rare situations, neural responses can serve as a reliable signal of preference, temperament, and even behavioral inclination.

Because many potentially aggressive dogs are not, in most situations, overtly aggressive, it may be difficult to behaviorally predict or assess risk until a potentially tragic incident has occurred, which may only occur upon exposure to a specific provocative stimulus or set of stimuli. As our previous work has shown, many dogs are able to inhibit a prepotent response, and they show a high degree of individual variability. This variability, though, can be significantly explained by the degree to which dogs deploy resources in the frontal lobe (Cook, Spivak & Berns, 2016). Overt aggressive behavior clearly depends on a range of interrelated factors, including situation, training, prior experience, capability for inhibition, and strength of aggressive drive. Moreover, the dog’s innate temperament and predilection for maintaining a relaxed emotional state versus an anxious emotional state that triggers concomitant sympathetic nervous system arousal is highly relevant to whether the dog will react aggressively upon exposure to a potentially provocative stimulus. Functional neuroimaging offers the chance to examine neural activations related to aggressive drive or inclination, even in the absence of overt aggressive behavior.

Building off of prior work by Harris & Prouvost (2014), we conducted an fMRI study in which dogs witnessed their primary caregiver provide a hot dog reward either to a realistic-looking fake dog, or a large red bucket. Brain response in an *a priori*, anatomically defined, amygdala region (Berns et al., 2017) was assessed in both conditions and used as an index of arousal. Activation levels were compared to temperament scores perviously extracted from C-BARQ assessments conducted by the dogs’ owners and found to correlate with measures of aggression.

## Materials and Methods

### Participants

Participants were 13 domestic dogs, and all dogs/owners were volunteers from the Atlanta area. All dogs had perviously completed one or more scans for the project and had demonstrated the ability to remain still during training and scanning (Berns, Spivak & Brooks, 2012; Cook, Spivak & Berns, 2014; Cook et al., 2016). This study was performed in accordance with the recommendations in the Guide for the Care and Use of Laboratory Animals of the National Institutes of Health. The study was approved by the Emory University IACUC (Protocol DAR-2002879-091817BA), and all owners gave written consent for their dog’s participation in the study. Because the dogs were already skilled in the MRI process, no additional training for the current study was required. As in our prior experiments (see Berns, Brooks & Spivak, 2013), for the purpose of hearing protection during live-scanning, all participants wore MuttMuffs® or ear plugs secured with vet wrap.

### Imaging

All scans were acquired on a 3T Siemens Trio MRI, and scan parameters were similar to those in Cook, Spivak & Berns (2014). Functional scans used a single-shot echo-planar imaging (EPI) sequence to acquire volumes of 22 sequential 2.5 mm slices with a 20% gap (TE=25 ms, TR = 1200 ms, flip angle = 70 degrees, 64×64 matrix, 3 mm in-plane voxel size, FOV=192 mm). A T2-weighted structural image was perviously acquired during one of our earlier experiments using a turbo spin-echo sequence (25-36 2 mm slices, TR = 3940 ms, TE = 8.9 ms, flip angle = 131 degrees, 26 echo trains, 128×128 matrix, FOV=192 mm). Three runs of up to 800 functional volumes were acquired for each dog, with each run lasting 12-16 minutes.

### Experimental Design

There were three trial types: 1) dog gets food; 2) fake-dog gets food; and 3) food is deposited in a bucket. Each trial began with the presentation of an object on the end of a wooden stick for 6-7 s. Then, depending on the trial type, the owner either fed the subject dog, put the food in a small, hidden pouch attached to the fake-dog’s mouth, or put the food in the bucket. The subject dog in the scanner was able to see these outcomes (Fig. 1). Each of the three runs contained 30 trials of each type in random order.

**Figure 1.**
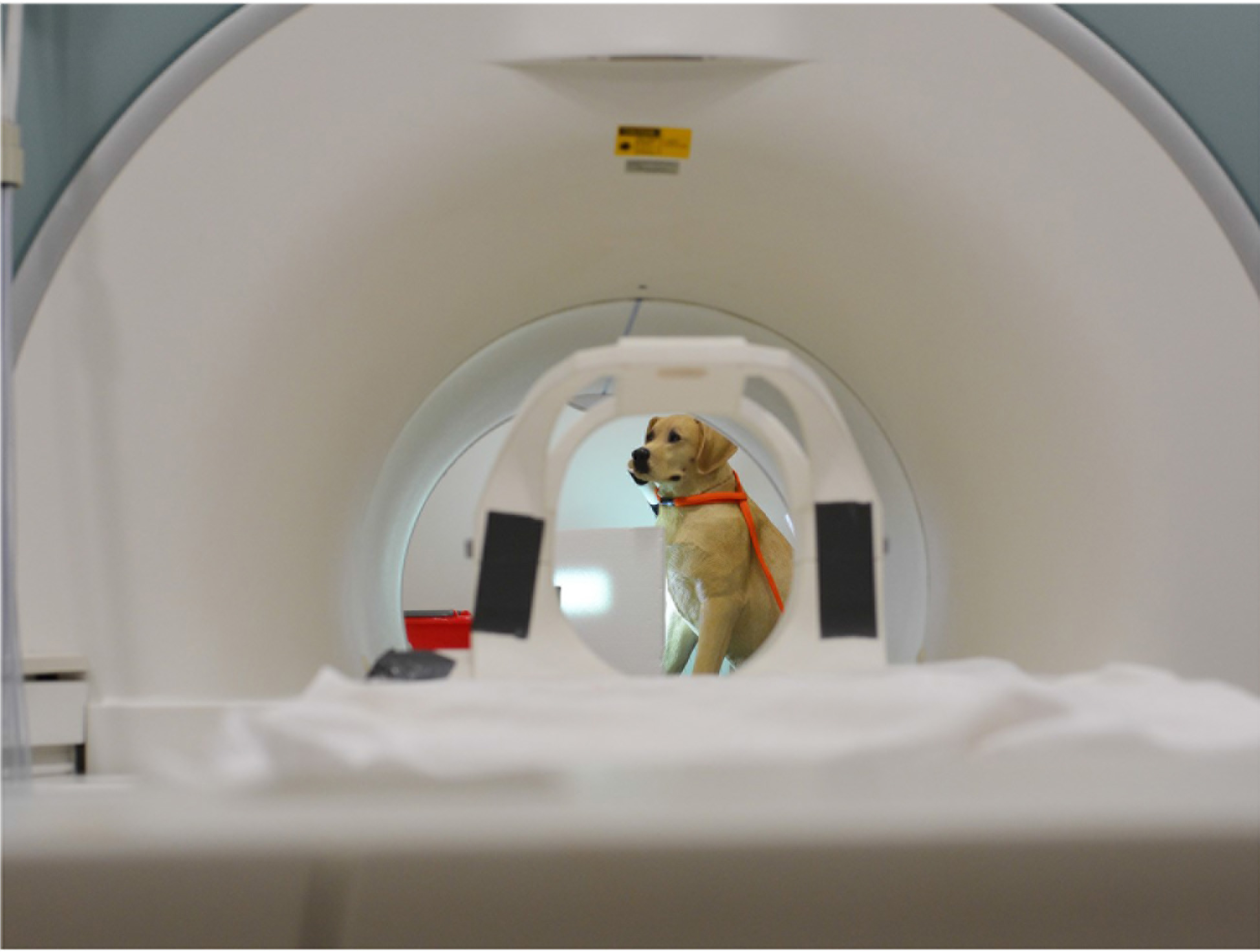
A view of the fake dog through the scanner bore. To “feed” the fake dog, the owner placed food in a tube behind the dog’s muzzle.

Trial events (onset and offset of object presentations) were recorded by an observer out-of-sight of the subject via a four-button MRI-compatible button-box. A computer running PsychoPy (Peirce, 2009) was connected to the button-box via usb port, and recorded both the button-box responses by the observer and scanner sequence pulses.

### Analsis

Data preprocessing included motion correction, censoring and normalization using AFNI (NIH) and its associated functions. Two-pass, six-parameter affine motion correction was used with a hand-selected reference volume for each dog. Aggressive censoring (i.e., removing bad volumes from the fMRI time sequence) was used because dogs can move between trials and when consuming rewards. Data were censored when estimated motion was greater than 1 mm displacement scan-to-scan and based on outlier voxel signal intensities. Smoothing, normalization, and motion correction parameters were identical to those described perviously (Cook et al. 2016). A high-resolution canine brain atlas (Datta et al., 2012) was used as the template space for individual spatial transformations. The Advanced Normalization Tools (ANTs) software was used to spatially normalize the statistical maps of the contrasts of interest to the template brain using affine and symmetric normalization (SyN) nornlinear transformations (Avants et al., 2011).

Each subject’s motion-corrected, censored, smoothed images were analyzed within a General Linear Model (GLM) for each voxel in the brain using 3dDeconvolve (part of the AFNI suite). Nuisance regressors included the six motion parameters. A constant and linear drift term were included for each run to account for baseline shifts between runs as well as slow drifts unrelated to the experiment. Task-related regressors were modeled using AFNI’s GAM function with 3 s duration and were as follows: (1) onset of trial cue; (2) feeding of fake-dog (the moment the food was placed in the pouch); (3) food in bucket. These were modeled separately for each run so that we could measure any habituation that occurred across runs. The outcome of feeding the subject dog was not modeled because those volumes were censored due to excessive motion during the consumption of the food.

To identify the neural response associated with a realistic dog receiving a treat, but controlling for the fact that the subject didn’t get the expected treat, we formed one contrast of interest: [fake dog – bucket]. In theory, this isolated the social saliency of the fake dog.

Because our main hypothesis centered on the role of the amygdala in social saliency, we extracted mean beta values for this contrast from the left and right amygdala after spatial normalization to the atlas (Fig.2). Using a mixed-effect model in SPSS (v. 23, IBM) with identity covariance structure and maximum-likelyihood estimation, we tested for the following fixed-effects: sqrt(run-1), CBARQ dog-directed aggression score, and interaction of sqrt(run-1) with dog-directed aggression score. The rationale for including run number was to control for any habituation effects that might occur from repeated presentations of a stimulus, and using sqrt(run-1) allowed for a presumed curvilinear relationship to run number. CBARQ is a validated assessment criteria for evaluating canine temperament and behavior. The inclusion of CBARQ dog-directed aggression tested for the relationship between this behavioral trait and amygdala reactivity to the saliency of the fake dog, while the interaction with run number controlled for potentially different rates of habituation with different temperaments.

## Results

The mixed-effect model showed a significant relationship of amygdala activity to dog-directed aggression score [t(76)=3.10, p=0.003] as well as the interaction with sqrt(run-1) [t(76)=−2.78, p=0.007] (Table 1). The negative coefficient of the interaction of sqrt(run-1) × dog-aggression indicated that dogs with higher aggression not only had a higher amygdala activation to the fake dog, but also had a greater amount of habituation over the 3 runs. Because activation and habituation were correlated, we also ran the analysis on run 1 only (Figure 2 & Table 2). This confirmed the relationship of amygdala activation to dog-aggression [t(26)=2.45, p=0.022].

**Table 1.** Estimates of fixed effects for amygdala and contrast: [fake dog – bucket].

| **Effect** | **Estimate** | **Std. Error** | **df** | **t** | **Sig.** |
| --- | --- | --- | --- | --- | --- |
| Intercept | -0.20 | 0.14 | 76 | -1.41 | 0.164 |
| Sqrt(Run-1) | 0.12 | 0.14 | 76 | 0.82 | 0.414 |
| Dog Directed Aggression | 0.48 | 0.16 | 76 | 3.10 | 0.003 |
| Sqrt(Run-1) x Dog Directed Aggr. | -0.44 | 0.16 | 76 | -2.78 | 0.007 |

**Table 2.** Estimates of fixed effects (Run 1 only) for amygdala and contrast: [fake dog – bucket].

| **Effect** | **Estimate** | **Std. Error** | **df** | **t** | **Sig.** |
| --- | --- | --- | --- | --- | --- |
| Intercept | -0.21 | 0.18 | 26 | -1.18 | 0.251 |
| Dog Directed Aggression | 0.50 | 0.20 | 26 | 2.45 | 0.022 |

**Figure 2.**
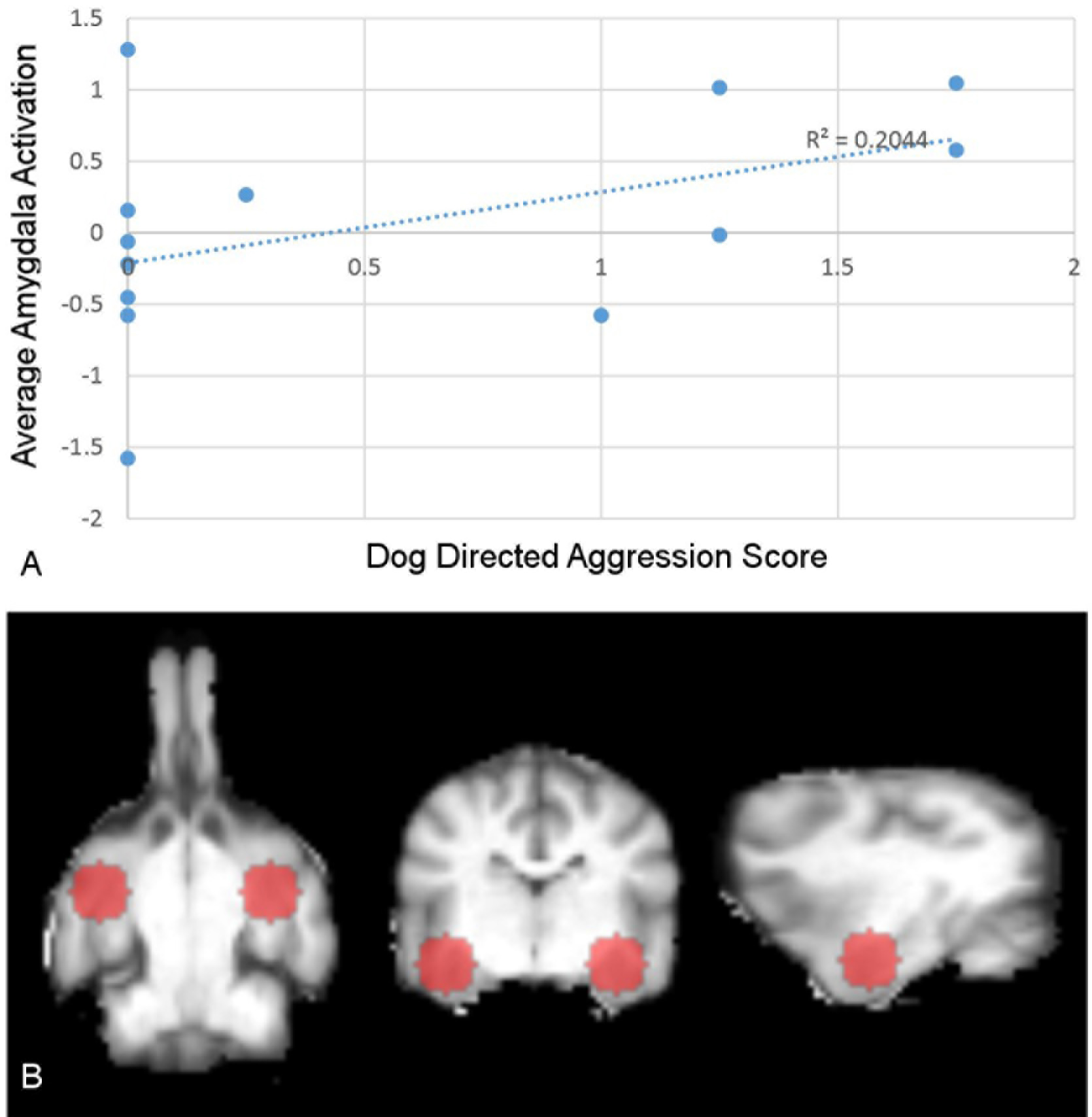
Relationship of amygdala activation to dog-directed aggression. *A)* Amygdala activation vs. dog-directed aggression score. Average amygdala activation during run 1 for each dog, plotted as a function of the dog-directed aggression score. There is a significant, positive correlation between the relative activation in the amygdala for [fake dog – bucket]. *B)* Anatomically defined, spherical, bilateral amygdala ROIs used to determine amygdala activation, shown in the transverse, coronal, and sagittal planes.

## Discussion

In 13 domestic dogs cooperatively scanned with awake-fMRI, aggressive temperament was positively correlated with bilateral amygdala activation when viewing their respective owners providing food to arealistic looking fake dog relative to dropping the food in a bucket. Because dogs were rewarded with food on one third of trials, both the bucket and dog conditions involved loss of potential reward, and were differentiated only by the end-point for those rewards. Across all dogs, bilateral amygdala activation to the dog versus bucket conditions habituated across three experimental runs of 30 trials each.

Dog-dog and dog-human aggression is troublingly common, and results can bes catastrophic (Overall & Love, 2001). Although folk theories are rampant, there is little prior scientific knowledge on the biological and neurobiological underpinnings of aggression in domestic dogs. Our findings suggest that aggressive dogs are likely to show increased arousal--as indexed by amygdala activation--when their owners interact with other dogs in a food context. Importantly, none of our subjects left the scanner or showed any overt signs of aggression when food was provided to the false dog during imaging. In addition, even our most aggressive dogs were well trained and relatively well-mannered in the context of the MRI. Together, these facts suggest that covert arousal can be increased in aggressive dogs in certain situations, even without overt behavioral manifestations. Actual aggressive behavior is likely to be preceded by covert arousal, and there may be covert arousal in many situations when no actual aggression occurs. Interaction of owner with another dog may be a dangerous trigger for aggression in certain dogs, even if, in most cases, aggression does not occur. As discussed in (Cook, Spivak & Berns, 2016), dogs with poor inhibitory control and high degrees of covert aggression might be most at risk.

We also measured significant habituation of the amygdala response across experimental sessions, but this was seen only in the aggressive dogs, who were the dogs who had amygdala activation in the first place. Notably, this activation was maximal in the first run but effectively nonexistent in the second and third runs. This suggests that behavioral interventions involving controlled exposure to interactions between owner and other dogs might be an effective therapeutic intervention for dogs prone to aggressive behavior in these contexts. It may also be that exposure-based interventions would be effective for dogs who show aggression in other contexts as well. Certainly there is a robust literature indicating the value of exposure therapy in humans with anxiety and other high-arousal disorders (Davis, 2002; Hofmann, 2008). It may also be that aggressive dogs would respond well to drugs used to treat high arousal in humans, such as beta-blockers and alpha2-agonists, some of which have been used clinically in this population (Dodman, 1998)

Although not conclusive, our findings may also be relevant to social resource guarding in dogs. Recent behavioral findings indicate that dogs show a tendency toward aggressive behaviors when their owners show affection toward a fake dog as opposed to a neutral inanimate object (Harris & Prouvost, 2014), and this has been likened to human jealousy, or “proto-jealousy.” Certainly the bond between owner and dog is central to the socioecology of domestic dogs, and a number of social species have been shown to be covetous of attention/access to conspecifics (Panksepp, 2010). In previous research, we have shown that some dogs show activation in the ventral caudate--associated with reward anticipation--to receipt of praise (Cook et al., 2016). Many domestic dogs highly value owner attention, and may desire to protect their access to it.

In addition to responding to a potential social threat, our dogs might also have been responding to the simple receipt of reward by a conspecific, regardless of the role the owner played in delivering it. As with primates, dogs have also shown some sensitivity to “reward inequity,” that is, they react with aversion when a conspecific receives greater reward than they do (Range et al., 2009). Although reward was balanced between subject and fake dog in our study, all dogs in the study were accustomed to being fed in the imaging context and may have viewed the false dog as an interloper. Importantly, whether something like proto-jealousy or reward inequity drove the observed amygdala response, there was a clear differential when the reward was deposited in a bucket, suggesting that our subjects were sensitive to the target of human attention/food reward, not just loss of a potential treat.

We could not rule out, in the current study, an alternative interpretation of our findings. It may be that aggressive dogs show increased amygdala activation an time the attend to a conspecific, regardless of context. Although the fake dog was present throughout the imaging sessions, and was always fully visible to the subjects, having the human deliver a food reward to the fake dog likely increased attentional focus on the fake dog. Future work might seek to disentangle this by finding alternate, non-social means to direct attention to the fake dog, or examining brain activation when the fake dog was visually accessible as opposed to hidden.

In addition, the complexities of the scanning environment necessitated our using a fake, instead of a real, dog. False dogs have also been used in behavioral studyies, and we observed apparently social reactions (e.g., growling, sniffing) to our fake dog in pilot work, but more ecologyically valid work with real conspecifics would be of value. Determining how to control the behavior of a real dog during testing so as to avoid experimental confounds is paramount. Furthermore, with real dogs the neurobiology of social status may be studyied, which might answer questions as to the effect of status and relationship dynamics upon the probability of affiliative behavior or jealousy and aggression.

Dog-dog, and dog-human aggression impact millions of people world-wide. Our findings highlyight the potential mediating factor of covert arousal, and the compounding roles of temperament and human attention. Moreover, the study and understanding of covert aggression potentially has highly valuable implications to pet owners and society as a whole. Pet owners often cite that their dog “gave no warning” prior to an attack. Yet, the onset of amygdala activation might be a cogent warning, Further study may determine a visible correlate derived from changes in facial countenance or body language that indicate amygdala activity. Importantly, our findings also suggest covert arousal may habituate with exposure. In addition to the behavior modification questions posed above, future work might seek to examine the neurobiological effects of behavioral and pharmacological interventions with aggressive dogs.

## Acknowledgments

Thank you to all of the owners who trained their dogs for this study: Lorrie Backer, Darlene Coyne, Vicki D’Amico, Diana Delatour, Marianne Ferraro, Cindy Keen, Patricia King, Cecilia Kurland, Claire Mancebo, Patricia Rudi, Lisa Tallant, and Nicole Zitron.

## Author Contributions

P.C., M.S, and G.B. designed the research; all authors collected data; P.C & G.B. analyzed data; all authors wrote the paper.

